# Bone conducted responses using the parallel auditory brainstem response (pABR) paradigm

**DOI:** 10.1101/2025.10.07.680974

**Authors:** Melissa J. Polonenko, Ross K. Maddox

## Abstract

**Objectives:** Auditory brainstem responses (ABRs) are used to diagnose hearing loss across multiple frequencies in infants and others who cannot participate in behavioral testing. The parallel ABR (pABR) paradigm measures response waveforms to all frequencies from 500– 8000 Hz in both ears all at once, rather than serially in one ear at a time. Work so far has been with air conducted stimuli through insert earphones, but bone conducted stimuli are also an essential part of diagnosing the type of hearing loss by determining the relative contributions of conductive and sensorineural hearing loss. This study aimed to confirm the feasibility of bone conduction pABR.

**Design:** We recruited a cohort of young adults with normal hearing. An intensity series of two-channel ABRs were measured to pABR stimuli played through ER-2 insert earphones as well as with a B-71 bone vibrator with contralateral masking. Behavioral thresholds were also measured to compare dB nHL correction factors. To keep sessions under 3.5 hours only one ear was tested with bone conduction. The ear with bone conduction and the order of transducers was counterbalanced across participants. Waveform morphologies for each transducer were compared by their wave V latencies and amplitudes and residual noise.

**Results:** For the same dB nHL levels, responses to both transducers aligned well and wave V amplitudes and latencies were similar. While stimulus artifacts can be an issue with bone conduction, the steps taken to mitigate it in air conduction pABR were also effective for bone conduction: clear bone conducted responses were measured and residual noise was similar for both transducers. Bone conducted responses to 8 kHz were weak, which was not surprising given the physics of bone conduction and the output capabilities of the bone vibrator. As expected, behavioral thresholds were higher for pABR than pure tone stimuli, and by the same offset for both transducers. The dB nHL values differed due to the difference in reference equivalent threshold levels for inserts and the bone vibrator.

**Conclusions:** Air and bone conduction provided similar waveforms, indicating that bone conduction pABR is feasible. Together with establishing dB nHL values, this work represents an important step towards translating the pABR for diagnosing all types of hearing loss.

## Introduction

The frequency-specific auditory brainstem response (ABR) is the primary method of diagnosing hearing loss in infants and others who cannot participate in behavioral testing. Thresholds at each frequency and in each ear are determined by varying level until a response is no longer detected. Air conduction (AC) thresholds using stimuli presented over earphones are sensitive to all types of hearing loss, such that they cannot discern sensorineural hearing loss from conductive or mixed losses. Using bone conduction (BC) transducers allows the stimuli to bypass the middle ear and stimulate the cochlea directly, making it sensitive only to sensorineural loss. Although tympanometry can indicate conductive middle ear issues, BC ABR is necessary to determine the relative contributions of conductive and sensorineural hearing loss. Because AC and BC thresholds can be interpreted together to determine the type of an individual’s hearing loss, BC is an essential part of ABR testing (Stapells & Oates 1997; Joint Committee on Infant Hearing 2019).

The parallel ABR (pABR) speeds up testing by measuring the responses to many frequencies in both ears all at once (Polonenko & Maddox 2019). The pABR quickly provides high quality responses that align with hearing thresholds in people with normal hearing (Polonenko & Maddox 2019; Polonenko & Maddox 2022), and ongoing work will test its accuracy in people with hearing loss. All work on the pABR so far has been with AC stimuli. While BC has been demonstrated to be effective and accurate with traditional ABR testing (Stapells & Oates 1997; Stapells 2000; Stapells 2011), the differences between traditional and pABR (specifically, the simultaneous stimulus presentation across frequencies and random timing) require that pABR be tested with BC as well.

The goal of this study is to provide proof of feasibility for using the pABR under BC. We recruited adults with normal hearing and recorded pABR using AC and BC transducers. We also determined BC correction factors through a behavioral experiment, as we did for AC in a previous study (Polonenko & Maddox 2022). The overall results confirm that using BC with the pABR presents no issues beyond those also faced in conventional serial ABR, answering a question that is important to the pABR’s translation to the clinic.

## Methods

### Participants

All research was performed under a single protocol approved by the University of Michigan IRBMED with a site-specific protocol approved by the University of Minnesota Institutional Review Board. All methods were in accord with the Declaration of Helsinki. Twenty-two adults aged 19 to 31 years (mean ± SD: 22.8 ± 2.7 years) were recruited and gave written consent prior to any testing. Participants were compensated for their time. There were 19 females, 2 males and 1 gender that is not exclusively male or female.

Wideband immittance, measured with an Interacoustics Titan, indicated normal middle ear function in all participants. Normal hearing thresholds were confirmed from 0.5 to 8 kHz using an Interacoustics AC40 audiometer coupled to IP3 earphones and a B-71 bone vibrator. The mean ± SD (range) pure-tone-average across the five octave frequencies was 4.9 ± 3.8 (1–12) dB HL for air-conduction (AC) and 1.9 ± 4.7 (-6–11) dB HL for bone-conduction (BC), with an air-bone-gap of 3.0 ± 3.9 (-2–12) dB.

Of these 22 participants, 2 did not complete the behavioral task at the time of the EEG and did not return for another session. One participant had behavioral data but unusable EEG data (the earplug was blocked or fell out during testing). Thus, one participant was recruited for only behavioral testing so that both the behavioral and EEG tasks had a total of 20 participants (with 18 participants having completed both tasks). To keep sessions under 3.5 hours, only one ear was tested with BC. Across the 20 participants, the ear tested with BC and the order of AC and BC testing were counterbalanced for each task.

### Stimuli

The pABR stimuli were constructed at a 48 kHz sampling rate as described previously (Polonenko & Maddox 2019). Briefly, 30 unique 1 s stereo tokens were made by: 1) creating Blackman-windowed 5-cycle tone pips centered at octave frequencies from 0.5 to 8 kHz; 2) separately creating a pulse train for each tone pip frequency by placing the tone pips randomly within a 1 s token and randomly inverting half the tone pips (akin to alternating phase); 3) summing the 5 random pulse trains together to create one channel, and 4) repeating the independent randomized process for the other channel. The number of tone pips in each token corresponded to the stimulus presentation rate. Each unique token was duplicated and then inverted in sequence to counter-phase the stimuli to further mitigate stimulus artifact for the EEG task. The sequence of tokens was repeated as many times as needed for the number of trials in the EEG task, whereas a token was randomly chosen with replacement for each presentation of the stimulus in the behavioral task. Details of this process can be found in Polonenko and Maddox (2019).

The tone pip stimuli were calibrated to a 70 dB peak-equivalent sound pressure level (peSPL) by matching the peak amplitudes of the tone pip to the amplitude of a 1 kHz pure tone that read 70 dB SPL on a Larson Davis Model 831 sound level meter when played through an ER-2 earphone attached to a 2-cc coupler. The other stimulus levels (L) were obtained by multiplying the reference tone pip by 10^(L – 70)/20^. Output of the bone vibrator was calibrated using a Brüel & Kjær 2203 Precision sound level meter connected to an artificial mastoid (re: dB dyne, 10 µN) and an ATTen 10621 oscilloscope (RMS voltage output).

For the behavioral task, pure tones were also created using 1 s sinusoids with amplitudes that were calibrated to the same 70 dB SPL tone. The pure tones were windowed with 35 ms raised cosine edges to match the audiometer settings used in the hearing screening, as done in our previous study (Polonenko & Maddox 2022).

### Stimulus presentation

Both the behavioral and EEG tasks used custom python scripts with the expyfun package (Larson et al. 2014) to present sound and triggers for the EEG through an RME digiface USB soundcard coupled with ER-2 insert earphones for AC and a B71 bone vibrator for BC testing. The same output voltages – calibrated to a 1 kHz pure tone via AC – were used for both transducers to allow easy switching between conditions and to determine the appropriate voltage corrections for future experiments that would seamlessly interchange between AC and BC testing. This choice means that the range of effective levels tested at the ear were different, which required post hoc alignment of responses, described below.

Because the main purpose of the study was proof-of-concept for parallel stimulation with BC testing, we decided to conduct a behavioral task to determine the dB nHL corrections in the same session as collecting EEG responses to ensure we had sufficient data for both tasks and could determine the BC levels. This resulted in some of BC EEG intensities having sensation levels (SLs) below threshold. Levels were corrected to 0 dB nHL and waveforms shifted down to match the closest 10-dB step-size for both transducers. Details are provided in the results section.

### Behavioral thresholds task

Behavioral thresholds were measured for pure tones (PT) and 20, 40, 60, and 80 stimuli/s rates to establish BC dB nHL reference values. We replicated the behavioral task from our previous study (Polonenko & Maddox 2022) that established the AC dB nHL correction factors. Briefly, participants sat in a darkened room and watched a computer screen containing a center white dot while pressing a button whenever they heard a sound. Stimuli were presented at a time randomly jittered between 1 to 4 s. Hearing thresholds were determined using an automated tracker based on the standard audiometric Hughson-Westlake procedure (Carhart & Jerger 1959) of 10 dB down / 5 dB up and a starting level of 40 dB (pe)SPL. If there was no response at the starting level, the level increased in 20 dB steps until there was a response or the level reached 85 dB (pe)SPL. Threshold was defined as the level at which 2 out of 3 responses were correct while the level ascended. pABR stimuli were presented in serial rather than parallel to determine thresholds for each tone pip frequency. All 25 threshold tracks were randomly presented over the course of the task (1 ear x 5 frequencies x 5 rates, with 0 Hz rate representing pure tones).

We identified an audio artifact after data had been collected that affected about half the participants’ AC thresholds. The artifact did not affect the BC results because of the higher voltage requirement to drive the bone vibrator. For this reason, AC behavioral data from this study were not used, with data from our previous study (Polonenko & Maddox 2022) used instead.

### EEG data collection

Participants reclined in a chair in a darkened audiobooth while 2-channel ABRs were recorded with EP-PreAmps connected to a BrainVision actiCHamp Plus amplifier. Passive Ag/AgCl electrodes were placed at Fz (active/non-inverting, split between both preamplifiers with a y-connector), Fpz (ground), and A1 and A2 (reference/inverting). Data were sampled at 10 kHz and high-pass filtered at 0.1 Hz. Triggers from the soundcard marked the beginning of each epoch to efficiently analyze each 1 s epoch of data in the frequency domain, as done before (Polonenko & Maddox 2019; Polonenko & Maddox 2022).

An intensity series was recorded to 6 intensities (ordered 20, 40, 60, 70, 50, 30 dB peSPL, voltage calibrated to an AC 1 kHz pure tone) of pABR stimuli at an average rate of 40 stimuli/s for 10 minutes each (600 x 1 second trials x 2 transducer conditions, total 120 minutes). This rate was previously determined as the most effective single rate for producing high-quality ABRs in a reasonable recording time (Polonenko & Maddox 2022). AC testing comprised dichotic stimuli. During BC testing, contralateral AC masking was presented at the same “level” (i.e., voltage output but not necessarily the same dB sensation level because of the different transducer frequency responses) but with 400 stimuli/s, which resulted in levels roughly 20 dB higher than at 40 stimuli/s.

The actual levels presented in the intensity series were calculated based on the calibrated levels in dB dyne for the voltage outputs of highest level (70 dB) and then adjusted based on the correction factors determined from the behavioral task to convert to dB nHL. This was done for both the AC (dB SPL to dB nHL) and BC (dB dyne to dB nHL) so that waveforms could be compared for the same levels. Due to the different voltage requirements to drive the two transducers, the BC levels were lower than the AC levels and the waveforms in the intensity series were shifted down to lower levels to best match those of the AC waveforms. Actual calculated dB nHL levels for the transducers across each frequency were calculated, with details given in the results section.

### Data analysis

#### Behavioral dB nHL values

The dB nHL reference values for BC were calculated as done before (Gorga et al. 1993; Stapells & Oates 1997; Sharma et al. 2003; Polonenko & Maddox 2022). To correct for the voltage outputs based on AC calibration, temporal integration of brief stimuli, as well as the participants’ own hearing sensitivity, the thresholds to pure tones were subtracted from the thresholds to the pABR stimuli. AC offsets from prior work (Polonenko & Maddox 2022) were used to compare whether offsets differed by transducer or if one set of correction factors could be used. A nested linear mixed effects regression was used to model these corrections, with a random intercept per participant by transducer and fixed effects of rate, logged frequency, transducer and their 2-way interactions and the 3-way interaction. The reference equivalent threshold vibratory force levels (RETVFLs) for the B-71 bone vibrator and the reference equivalent threshold sound pressure levels (RETSPLs) for the ER-2 inserts were added to the modeled corrections to finally give the reference values for 0 dB nHL.

#### ABR derivation

ABRs were derived using the *mne* package (Gramfort et al. 2013; Larson et al. 2023) as described in detail elsewhere (Polonenko & Maddox 2019; Polonenko & Maddox 2022). Briefly, the raw EEG was band-pass filtered from 30 to 1500 Hz using a first order Butterworth filter and then notch-filtered at odd multiples of 60 Hz to remove electrical line noise. The data were then epoched from 500 ms before the onset of each 1 s trial (demarked by the trigger) to 500 ms after, for a total of 2 s per epoch. Because of the random processes used to create the stimuli, this same EEG was used to derive waveforms to each of the tone pip frequencies in each ear. This was accomplished by first rectifying the impulse sequences used to create the random pulse trains and zero-padding them with 500 ms on each end to give 2 s impulse trains that matched the EEG epoch. The ABR for each epoch was then calculated as the circular cross-correlation of the 2 s rectified impulse trains and EEG epochs, normalized to the average number of pulses in the sequence (i.e. stimulation rate), and all performed in the frequency domain for efficiency. Due to the circular nature of the cross-correlation, the middle 1 s of the response was discarded and the last 500 ms was concatenated with the first 500 ms to give a final response from [-500, 500] ms, where 0 ms denotes the onset of the tone pip. This was repeated for each of 10 tone pips and for each of the two EEG channels for every epoch. Averaged ABRs were created by weighting each epoch by the inverse of its variance relative to the summed inverse variances of all epochs for that condition so that noisy trials contributed less to the average.

#### Wave V peaks

Responses to bone- and air-conducted stimuli were compared by characterizing wave V peak amplitude and latency. Wave V peaks were manually picked by the first author at two different intervals, 4 weeks apart. Manual picks were done to mimic what is done in the clinic and to avoid spurious peaks being chosen due to noise, especially for the broader low frequency tone pips and for smaller responses at levels closer to threshold. Peaks were picked for the AC and BC responses averaged across channels, and for the BC responses in the ipsilateral and contralateral channels. Picks were repeatable across frequencies and types of stimuli, with intraclass correlation for absolute agreement (ICC3) coefficients > 0.994 [latency: *F*(4, 4) = 2367.0, *p* < 0.001, *ICC3* = 0.999, 95% CI = (0.99, 1.00); amplitude: *F*(4, 4) = 377.7, *p* < 0.001, *ICC3* = 0.995, 95% CI = 0.995 (0.95, 1.00)].

#### Statistical analysis

The primary goal was to determine if pABR with BC is feasible. To achieve this, we compared behavioral data and ABR morphology between AC and BC using linear mixed effects regression (LMER). EEG residual noise was analyzed with independent *t*-tests (mu = 0) and p-values were corrected for family-wise error using False Discovery Rate (FDR; Benjamini & Hochberg 1995). All models and their power analyses were performed using the *lmerTest* and *simR* packages in *R* (Kuznetsova et al. 2017; Green & MacLeod 2016; R Core Team 2025). Power was calculated using a likelihood ratio test performed on 1000 Monte Carlo permutations of the response variables based on the fitted models. The ICC3 coefficients were calculated with *pingouin* (Vallat 2018) package in python.

All data analysis and visualization besides the linear mixed effects modeling was done in python with custom scripts with support from *mne* (Gramfort et al. 2013) for the EEG data analysis pipelines.

## Results

### Behavioral thresholds for BC pABR normative values

Behavioral thresholds for BC stimuli are shown in Figure 1. Behavioral thresholds for the same amplifier output were higher for pABR than pure tone stimuli. These offsets are expected due to the temporal integration of the brief stimuli, and consistent with prior work with AC pABR stimuli (Polonenko & Maddox 2022). Also consistent with prior work, mean offsets were between 10 and 22 dB, decreased with stimulation rate, and differed by frequency.

**Figure 1.**
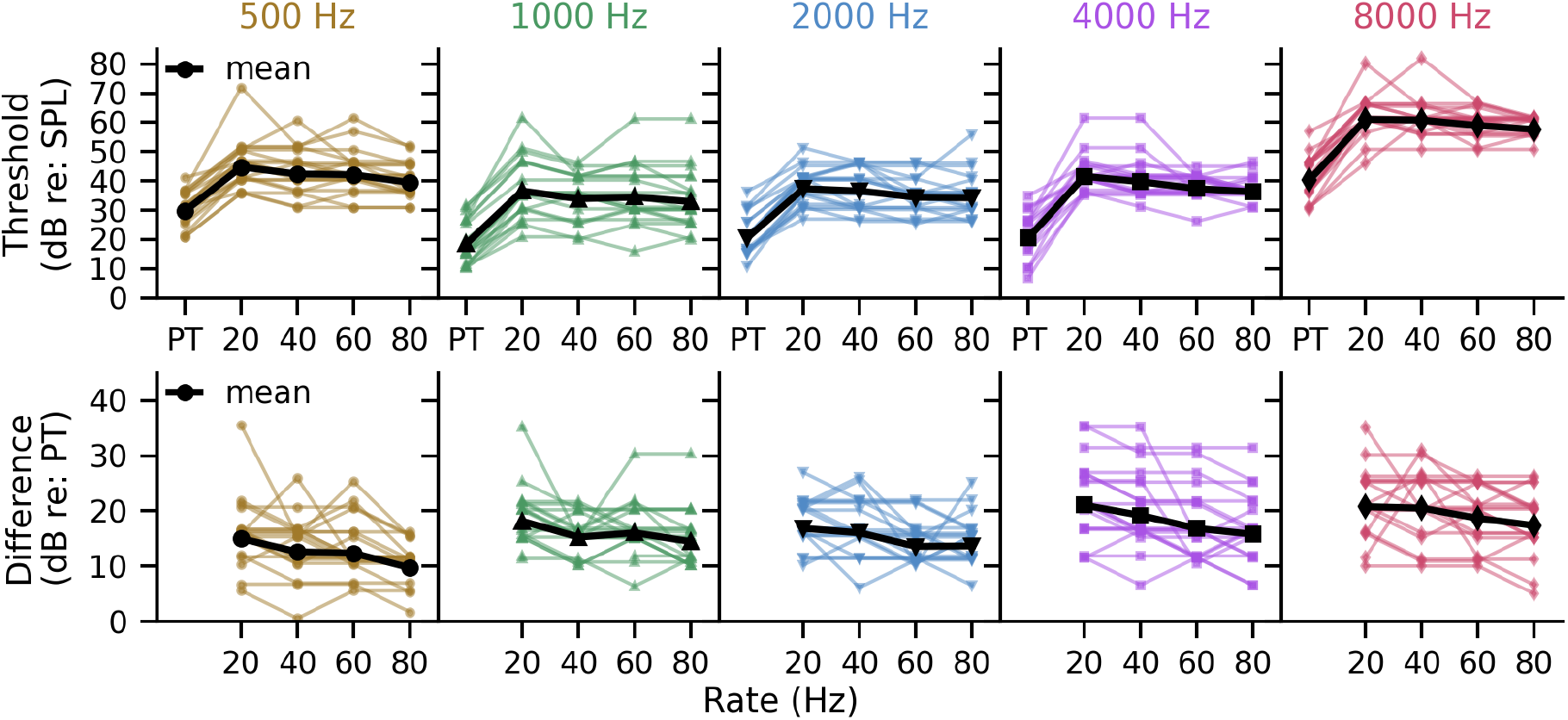
Bone conduction (A) thresholds and (B) offsets of the pABR stimuli compared to pure tones (PT) thresholds. Thin colored lines denote individual data and black thick lines the mean.

To determine the correction factors for pABR stimuli, we fit a nested linear mixed effects model to the behavioral offsets. We used these BC offsets and AC offsets determined in our prior study (Polonenko & Maddox 2022). The model included fixed effects of stimulus rate, logged frequency, transducer and their interactions, along with a random intercept for each participant by transducer (Table 1A). The model fit can be seen in Figure 2A as dark solid lines for BC and light dashed lines for AC. The BC offsets were similar to the AC offsets (all *p* > 0.407, Table 1A; modeled correction factors for both AC and BC are found in Supplemental Digital Content, Supplemental Table 1) and they matched at the standard stimulus rate of 40 stimuli/s, which was used in the EEG task.

**Table 1.** Linear mixed effects models of correction factors for pABR stimuli.

| Fixed Effect | Estimate | SE | df | t | p | Power<br>(95% CI) |
| --- | --- | --- | --- | --- | --- | --- |
| A) Full model (CF ~ Rate * LogFreq * Trd + (1 \| Participant:Trd)) |  |  |  |  |  |  |
| Intercept (AC) | 5.399 | 4.121 | 1114.5 | 1.310 | 0.190 | 0.274<br>(0.247, 0.303) |
| Rate | -0.176 | 0.075 | 1099.0 | -2.361 | 0.018 | 0.673<br>(0.643, 0.702) |
| LogFreq | 4.118 | 1.233 | 1099.2 | 3.341 | <0.001 | 0.936<br>(0.919, 0.950) |
| Trd (BC) | -3.399 | 7.184 | 1114.0 | -0.473 | 0.636 | 0.069<br>(0.054, 0.087) |
| Rate:LogFreq | 0.036 | 0.022 | 1098.9 | 1.630 | 0.103 | 0.355<br>(0.325, 0.386) |
| Rate:Trd(BC) | 0.108 | 0.131 | 1098.3 | 0.829 | 0.407 | 0.136<br>(0.115, 0.159) |
| LogFreq:Trd (BC) | 0.221 | 2.152 | 1098.3 | 0.568 | 0.570 | 0.066<br>(0.051, 0.083) |
| Rate:LogFreq:Trd(BC) | -0.037 | 0.039 | 1098.2 | -0.934 | 0.351 | 0.152<br>(0.130, 0.176) |
| B) Parsimonious model (CF ~ Rate * LogFreq + (1 \| Participant:Trd)) |  |  |  |  |  |  |
| Intercept | 4.306 | 3.374 | 1117.1 | 1.276 | 0.202 | 0.249<br>(0.222, 0.277) |
| Rate | -0.141 | 0.061 | 1101.7 | -2.307 | 0.021 | 0.628<br>(0.597, 0.658) |
| LogFreq | 4.512 | 1.010 | 1101.8 | 4.467 | <0.001 | 0.988<br>(0.979, 0.994) |
| Rate:LogFreq | 0.025 | 0.018 | 1101.6 | 1.343 | 0.180 | 0.281<br>(0.253, 0.310) |

Note: AC data from (XX REF 2021); LogFreq = $\log_{10}$ (frequency in Hz), Trd = transducer, CF = correction factor, AC = air conduction (default in the full model), BC = bone conduction, CI = confidence interval, SE = standard error

**Figure 2.**
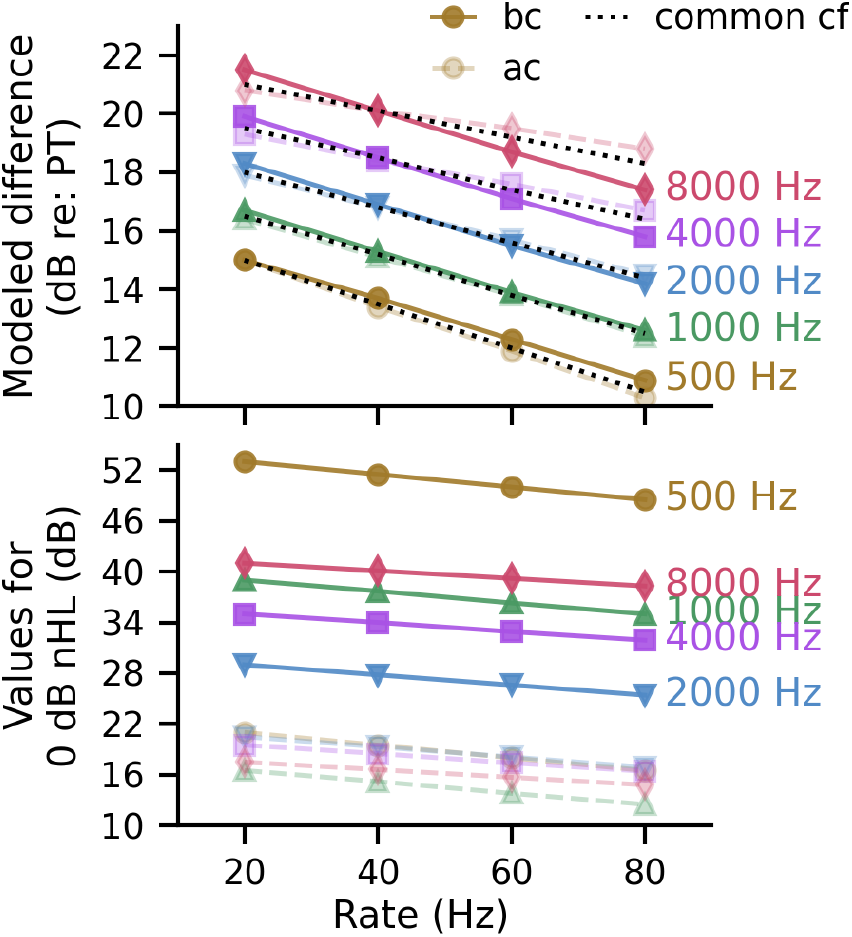
Modeled (A) offsets for bone- (solid line) and air- (dashed line) conduction and their combination (black dotted line). (B) values for 0 dB nHL, calculated as the modeled difference plus the RETVFL (BC) or RETSPL (AC).

Thus, we ran a reduced, more parsimonious model that could explain the data and give one set of modeled correction factors regardless of transducer, given in Table 1B and shown as the black dotted lines in Figure 2A.

BC and AC reference levels differ, and thus so do voltage output requirements between the two transducers, with the BC transducer requiring higher input levels to exert the same force on the ear. We computed the reference levels for 0 dB nHL by using the common set of modeled correction factors and adding the B-71 RETVFL for BC and the ER-2 RETSPL for AC, which are shown in Figure 2B. The large variation in RETVFL results in the AC values being somewhat visually compressed in the figure. The calibration values underlying these plots can be found in Table 2.

**Table 2.** Correction and normative values for converting dB SPL or VFL to 0 dB nHL.

| Rate | Tone pip |  |  |  |  |
| --- | --- | --- | --- | --- | --- |
|  | 500 Hz | 1000 Hz | 2000 Hz | 4000 Hz | 8000 Hz |
| A) Modeled correction factors (dB) [CF] |  |  |  |  |  |
| 20 Hz | 15.0 | 16.5 | 18.0 | 19.5 | 21.0 |
| 40 Hz | 13.5 | 15.2 | 16.8 | 18.5 | 20.1 |
| 60 Hz | 12.0 | 13.8 | 15.6 | 17.4 | 19.2 |
| 80 Hz | 10.5 | 12.5 | 14.4 | 16.4 | 18.3 |
| B) Reference equivalent thresholds (RET) for the transducers (dB) |  |  |  |  |  |
| RETVFL (BC) | 38.0 | 22.5 | 11.0 | 15.5 | 20.0 |
| RETSPL (AC) | 6.0 | 0.0 | 2.5 | 0.0 | -3.5 |
| C) BC (B-71) normative values for 0 dB nHL (dB) [CF + RETVFL] |  |  |  |  |  |
| 20 Hz | 53.0 | 39.0 | 29.0 | 35.0 | 41.0 |
| 40 Hz | 51.5 | 37.7 | 27.8 | 34.0 | 40.1 |
| 60 Hz | 50.0 | 36.3 | 26.6 | 32.9 | 39.2 |
| 80 Hz | 48.5 | 35.0 | 25.4 | 31.9 | 38.3 |
| D) AC (ER-2) normative values for 0 dB nHL (dB) [CF + RETSPL] |  |  |  |  |  |
| 20 Hz | 21.0 | 16.5 | 20.5 | 19.5 | 17.5 |
| 40 Hz | 19.5 | 15.2 | 19.3 | 18.5 | 16.6 |
| 60 Hz | 18.0 | 13.8 | 18.1 | 17.4 | 15.7 |
| 80 Hz | 16.5 | 12.5 | 16.9 | 16.4 | 14.8 |

Note: AC values from (REF 2021); VFL = vibratory force level; SPL = sound pressure level, CF = correction factor, AC = air conduction, BC = bone conduction

### Levels for the EEG data

Sound played with the same output voltages produced higher thresholds for BC than AC, due to differences in the transducers and the amount of energy reaching the cochlea. With the same voltage output, BC stimuli had lower levels as measured with a sound level meter coupled to an artificial mastoid (BC) or 2cc coupler (AC). The levels for voltage outputs of the 70 dB voltage outputs were converted to dB nHL using the measured dB dyne or SPL and subtracting the appropriate reference values for 0 dB nHL in Table 2C&D. Thus, the loudest 10-dB step-size level for the EEG data was ∼50 dB nHL for AC (50.5, 54.8, 50.7, 51.5, 53.4 dB nHL for 500 through 8000 Hz respectively) and ∼30 dB nHL for BC (29.8, 36.3, 38.4, 26.0, 20.4 dB nHL for 500 through 8000 Hz respectively), resulting in a difference between BC and AC levels of 20.8, 18.5, 12.3, 25.5, and 33.0 dB for 500 through 8000 Hz respectively. To compare waveforms for similar levels, the BC levels were shifted down by these calculated differences rounded to the nearest multiple of 10. For 4000 Hz, the AC-BC difference of 25.5 dB was in between 10 dB steps and we chose -20 instead of -30 because the waveforms aligned better. This resulted in the BC waveforms having a level -5.5 dB instead of +4.5 dB relative to the AC waveforms. Thus, the corrections were: 20, 20, 10, 20, and 30 dB by frequency respectively. In those plots (Figure 3), this quantized shift resulted in the BC waveforms having levels that were off from the AC levels by -0.8, +1.5, -2.3, -5.5, and -3 dB across frequency from 500 through 8000 Hz respectively. That error was not present in the horizontal axes of Figure 4, where the non-quantized (i.e., actual) offsets were used to compare wave V peak amplitudes and latencies. These calculated offsets are represented in Supplemental Digital Content, Supplemental Table 2.

**Figure 3.**
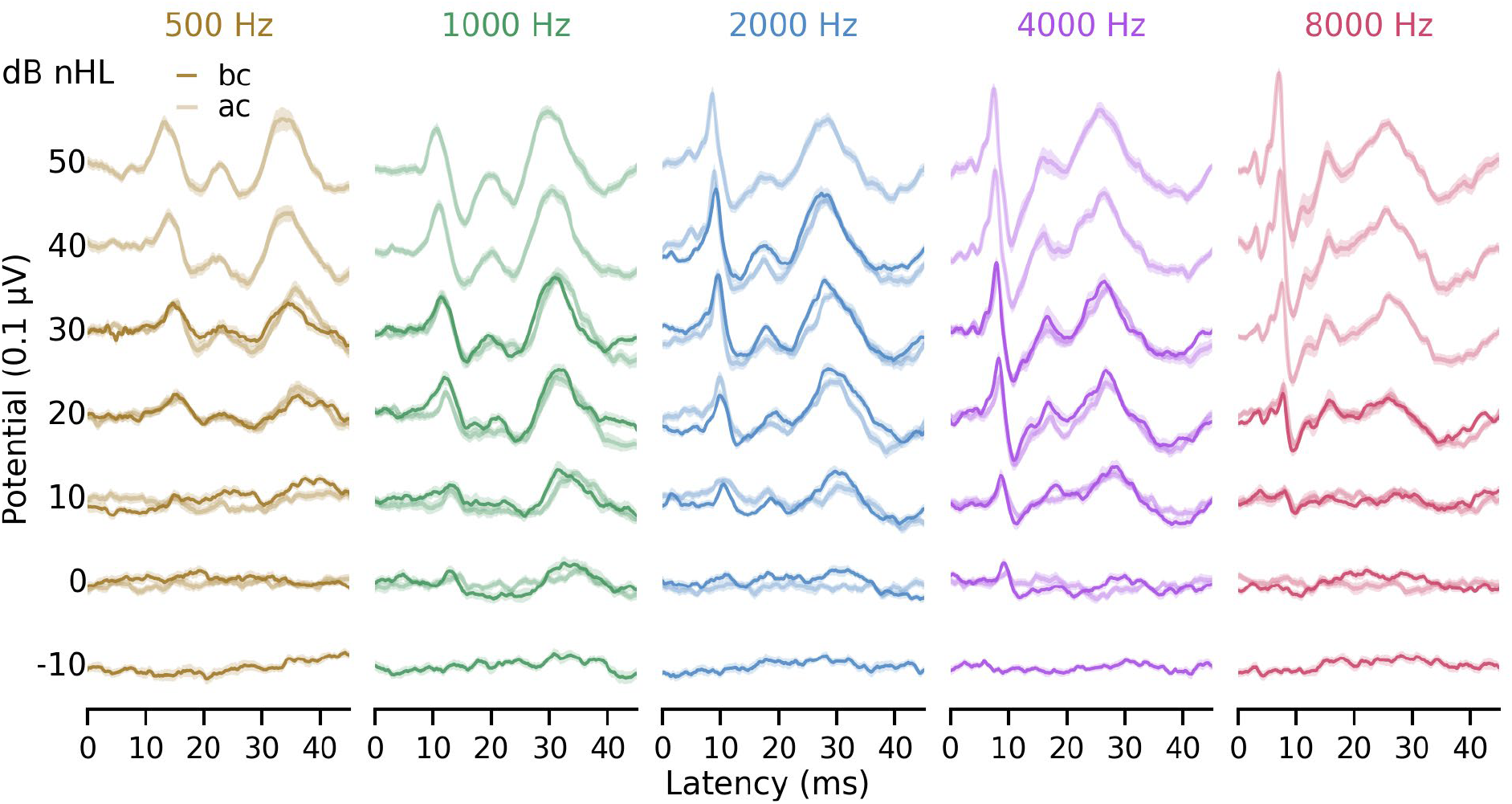
Waveforms for bone- (BC, dark thin line) and air- (AC, light thick line) conduction at levels matched to the nearest 10-dB step size.

**Figure 4.**
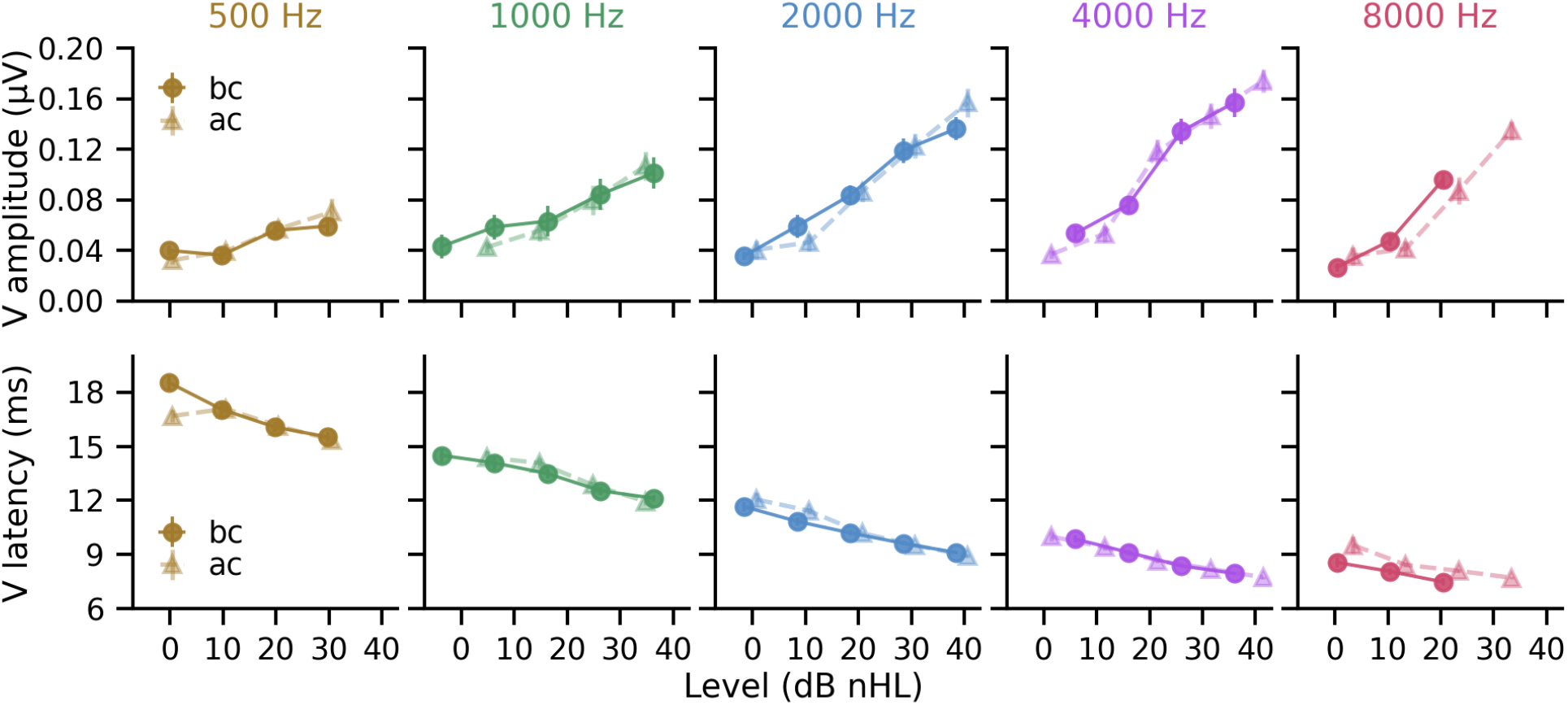
Wave V peak amplitude (top) and latency (bottom) for bone- (BC, circles and solid lines) and air- (AC, triangles and dashed lines) conducted stimuli at their actual dB nHL levels. Data is shown if there are at least 4 participants with data for that level. Error bars represent ±SE when visible.

### pABR responses are similar for bone and air conduction

Overall, waveforms were similar for bone- and air-conducted stimuli at similar levels, shown in Figure 3. There were some minor differences in the waveforms, which could be related to different runs in the experiment, but also the offsets to plot the waveforms together in 10 dB levels. In general, the BC waveforms were clear and qualitatively did not appear much noisier than the AC waveforms: the steps we took to mitigate BC artifact were effective. Furthermore, residual noise was similar for both AC and BC (linear mixed effects model all *p* > 0.486, shown in Supplemental Digital Content, Supplemental Figure 1 and Supplemental Table 3), with an overall mean ± SE of 25.14 ± 0.81 nV for BC and 25.47 ± 1.15 nV for AC. Robust responses were evoked for most tone pips, except for 8000 Hz at levels higher than 20 dB nHL. However, the voltage outputs used in this study were not sufficient to drive the B-71 vibrator at levels beyond ∼ 20 dB nHL for 8000 Hz, which is also a constraint for traditional ABR systems. Thus, while plotted, the 8000 Hz data were not included in the analysis.

To investigate the waveforms further, wave V peak amplitudes and latencies were compared using the calibrated levels for the two transducers, shown in Figure 4. We used linear mixed effects modeling, with fixed effects of level, logged frequency, transducer and their interactions, along with a random intercept for each participant. Levels beyond -10 to 45 dB were excluded to ensure data included responses above threshold and from at least 4 of the 20 participants at each level. Details of the models for amplitude and latency are given in Table 3A and B respectively. As seen in prior work (Polonenko & Maddox 2019), wave V amplitudes increased with increasing level and with increasing frequency (all *p* < 0.001), but BC stimuli did not change these relationships (intercept for BC and all its interactions had *p* > 0.257). BC also did not affect wave V latencies (BC intercept, 2-way and 3-way interactions, all *p* > 0.767), which showed the typical decrease with increasing frequency and increasing level (all *p* < 0.008).

**Table 3.** Linear mixed effects models of wave V peaks for pABR stimuli.

| Fixed Effect | Estimate | SE | df | t | p | Power<br>(95% CI) |
| --- | --- | --- | --- | --- | --- | --- |
| A) Amplitude (nV ~ dBs * LogFreq * Trd + (1 \| Participant)) |  |  |  |  |  |  |
| Intercept | -6.643 | 37.766 | 661.5 | -0.176 | 0.860 | 0.071<br>(0.056, 0.089) |
| dBs | -4.462 | 1.329 | 644.7 | -3.357 | <0.001 | 0.928<br>(0.910, 0.943) |
| LogFreq | 8.361 | 11.513 | 645.0 | 0.726 | 0.470 | 0.121<br>(0.101, 0.143) |
| Trd(BC) | 40.766 | 48.811 | 644.5 | 0.835 | 0.404 | 0.156<br>(0.134, 0.180) |
| dBs:LogFreq | 2.305 | 0.413 | 644.6 | 5.587 | <0.001 | 1.000<br>(0.996, 1.000) |
| dBs:Trd(BC) | -2.248 | 1.981 | 644.3 | -1.135 | 0.257 | 0.210<br>(0.185, 0.237) |
| LogFreq:Trd(BC) | -8.902 | 15.064 | 644.5 | -0.591 | 0.555 | 0.094<br>(0.077, 0.114) |
| dBs:LogFreq:Trd(BC) | 0.554 | 0.612 | 644.3 | 0.904 | 0.366 | 0.143<br>(0.122, 0.166) |
| B) Latency (ms ~ dBs * LogFreq * Trd + (1 \| Participant)) |  |  |  |  |  |  |
| Intercept | 41.262 | 1.046 | 656.5 | 39.438 | <0.001 | 1.000<br>(0.996, 1.000) |
| dBs | -0.174 | 0.037 | 645.6 | -4.705 | <0.001 | 0.998<br>(0.993, 1.000) |
| LogFreq | -8.725 | 0.320 | 646.2 | -27.220 | <0.001 | 1.000<br>(0.996, 1.000) |
| Trd(BC) | -0.154 | 1.359 | 645.2 | -0.114 | 0.909 | 0.053<br>(0.040, 0.069) |
| dBs:LogFreq | 0.030 | 0.011 | 645.4 | 2.647 | 0.008 | 0.757<br>(0.729, 0.783) |
| dBs:Trd(BC) | 0.016 | 0.055 | 644.7 | 0.297 | 0.767 | 0.055<br>(0.042, 0.071) |
| LogFreq:Trd(BC) | 0.001 | 0.419 | 645.2 | 0.004 | 0.997 | 0.045<br>(0.033, 0.060) |
| dBs:LogFreq:Trd(BC) | -0.004 | 0.017 | 644.7 | -0.248 | 0.804 | 0.064<br>(0.050, 0.081) |

Note: filtered dBs within -10 to 45 (not inclusive) and removed 8 kHz
LogFreq = log<sub>10</sub>(frequency in Hz), Trd = transducer, BC = bone conduction, default in the full model is air conduction (AC); SE = standard error, CI = confidence interval

### BC responses between ipsilateral and contralateral channels

So far, we have shown responses averaged across channel to increase the signal-to-noise ratio. However, in clinical exams the ipsilateral versus contralateral channels are often compared for BC stimuli to determine the side that is most likely responding given the lack of interaural attenuation. The channel with the larger and earlier wave V is likely the side generating the response (Stapells & Ruben 1989; Foxe & Stapells 1993; Hatton et al. 2022; Ontario Ministry of Children and Youth Services 2020). We presented contralateral AC masking to help mitigate the potential stimulus cross-over, but compare the ipsilateral and contralateral responses for ∼20 dB nHL level in Figure 5. This level is the lowest with responses from most participants for all 5 tone pips and reflects the level that would be used to determine whether the BC thresholds are within the normal range. Overall, the wave V responses appeared smaller for the contralateral channel and either peaked at the same or later latency. While these changes suggest the ear ipsilateral to the bone vibrator is responding, these small changes across channel were only significantly different from zero for 500, 2000 and 4000 Hz for amplitude and for 1000 and 2000 Hz for latency (see Table 4; independent *t*-test *p* corrected with FDR).

**Table 4.** Independent t-tests of differences between ipsilateral and contralateral channels.

| Frequency | Estimate | 95% CI | df | t | P | p-adjusted* |
| --- | --- | --- | --- | --- | --- | --- |
| A) Amplitude (nV) |  |  |  |  |  |  |
| 500 Hz | 19.8 | 6.9, 32.8 | 18 | 3.228 | 0.0047 | 0.0155 |
| 1000 Hz | 17.2 | -2.2, 36.9 | 14 | 1.871 | 0.0824 | 0.1294 |
| 2000 Hz | 43.2 | 27.9, 58.5 | 18 | 5.937 | 0.00001 | 0.0001 |
| 4000 Hz | 23.7 | 4.7, 42.7 | 19 | 2.612 | 0.0171 | 0.0342 |
| 8000 Hz | 9.5 | -2.1, 21.1 | 18 | 1.715 | 0.1035 | 0.1294 |
| B) Latency (ms) |  |  |  |  |  |  |
| 500 Hz | -0.097 | -0.584, 0.389 | 18 | -0.420 | 0.6791 | 0.6791 |
| 1000 Hz | -1.277 | -2.221, -0.332 | 14 | -2.900 | 0.0116 | 0.0291 |
| 2000 Hz | -0.389 | -0.625, -0.153 | 18 | -3.468 | 0.0027 | 0.0137 |
| 4000 Hz | -0.107 | -0.425, 0.210 | 19 | -0.709 | 0.4868 | 0.5409 |
| 8000 Hz | -0.092 | -0.205, 0.021 | 18 | -1.715 | 0.1035 | 0.1294 |

Note: difference = ipsilateral – contralateral; \*adjusted with false discovery rate (FDR); CI = confidence interval, $\mu = 0$

**Figure 5.**
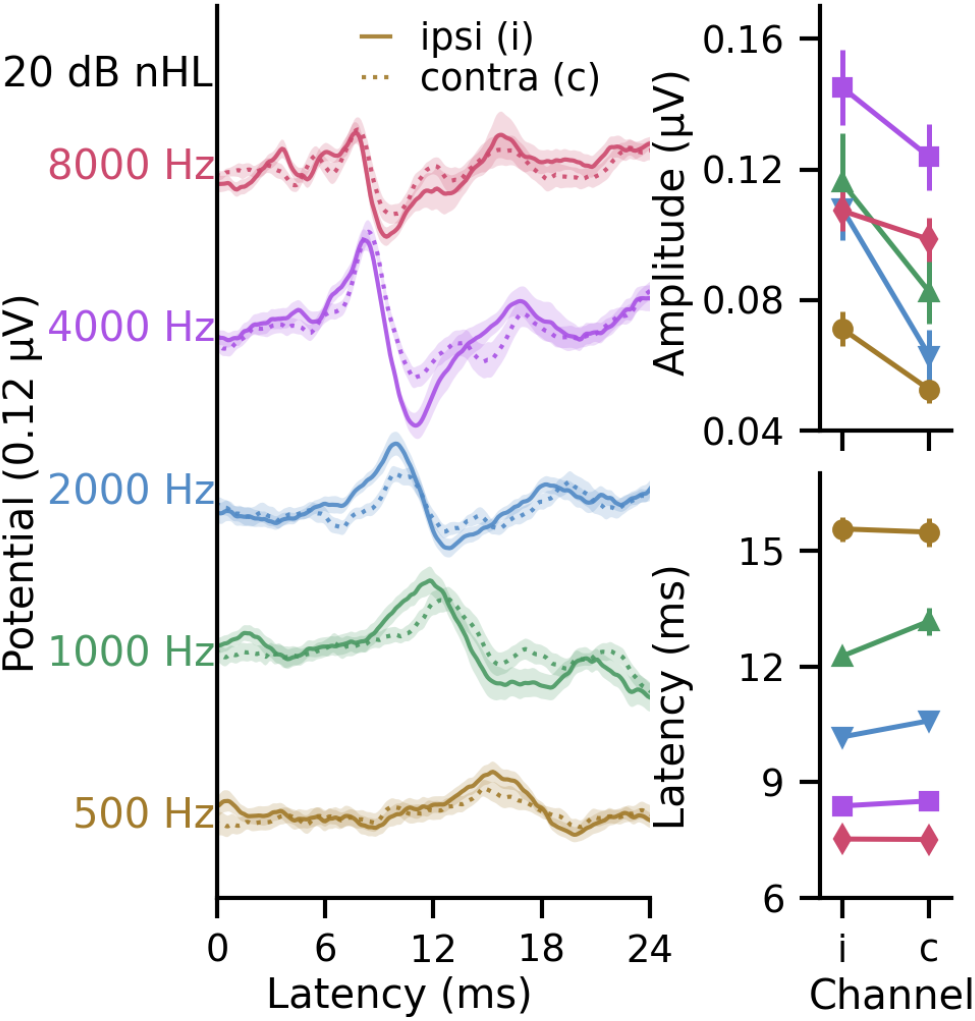
Bone-conducted waveforms (left) and wave V peak amplitudes and latencies (right) for the ipsilateral (ipsi, i; solid line) and contralateral (contra, c; dotted line) channels at levels ∼20 dB nHL (actual levels differ slightly, see text). Colors and symbols represent the different frequencies: 500 Hz (yellow circle), 1000 Hz (green triangle), 2000 Hz (blue inverted triangle), 4000 Hz (purple square), 8000 Hz (pink diamond).t

## Discussion

Here, we describe the pABR with air- (AC) and bone- (BC) conduction and establish one common set of correction values for both AC and BC transducers based on perceptual thresholds. These correction values are added to the RETSPL for AC and RETVFL for BC to give the reference values for dB nHL, which are higher for BC given the different voltage requirements to drive the bone vibrator. BC pABR produced robust responses with similar morphology and wave V peaks to AC at the same dB nHL levels. Using contralateral AC masking at the same voltage output as the BC but with 10x the stimulus rate (400 vs 40 stimulus/s) seemed to evoke responses from the ear ipsilateral to the bone vibrator placement. Thus, overall, the parallel stimulation of pABR is compatible with BC testing.

BC produced waveforms with similar morphology to AC with pABR. Very minimal artifact was observed, indicating that efforts to mitigate artifact were successful. Averaging epochs to pulse trains with half the pulses inverted, akin to alternating polarity, and the counter-phased presentation of epochs helped average out most, if not all, resonance from the bone vibrator. Furthermore, replications comprised of random subsets of pulses within the 1 s epochs gave more artifact-free averages than replications comprised of condensation and rarefaction, which was particularly helpful for the lower frequency tone pip responses that had longer stimuli that would “ring” closer in time with wave V.

When matched for level, BC pABR gave similarly sized responses across a range of intensities. This is expected given that the auditory system is stimulated with a similar level, once accounting for the transducer voltage outputs. Responses could be generated up to ∼30 dB nHL using similar voltage outputs as AC. This level delineates normal hearing from elevated thresholds and thus conductive from mixed or sensorineural hearing loss. A limitation of this study was the lower levels used for BC due to the necessity of using similar voltage outputs, as well as the frequency response of the B-71 oscillator. Now that the calibration levels are known for BC pABR, future studies could evaluate higher levels by having a different set of voltages for BC than AC while still using a combined paradigm. Such a paradigm would then better estimate elevated BC thresholds to delineate mixed from sensorineural hearing loss, although would still be limited by the B-71 vibrator’s physical capabilities.

Although responses matched well for AC and BC, BC was not as effective in evoking response at 8000 Hz. The B-71 is difficult to drive with enough force at 8000 Hz to test with higher levels, which is a constraint of the transducer and also a limitation to current clinical protocols. With the voltage outputs used in this study, high-frequency responses were recorded at 20 dB nHL in normal hearing adults. The presence of such responses could establish a normal hearing threshold but much higher levels, which may not be possible, would be needed to determine degree of any sensorineural component to a hearing loss at this tone pip frequency. Current protocols do not use 8000 Hz (Hatton et al. 2022; Joint Committee on Infant Hearing 2019; Ontario Ministry of Children and Youth Services 2020) and the BC pABR paradigm could remove 8000 Hz from the stimulus set so that high levels of an ineffective stimulus would be removed.

Our recording paradigm allows us to view the responses averaged across channels as well as the responses separately from each channel. In clinic, the ipsilateral and contralateral channels are compared to determine which ear is likely responding (Ontario Ministry of Children and Youth Services 2020; Hatton et al. 2022), which is the side/channel that has a response with the earlier and larger peak (Foxe & Stapells 1993; Stapells & Ruben 1989; Hatton et al. 2012). In infants this can often be done without contralateral masking due to some/higher interaural attenuation resulting from the lack of fused skull bones (Vander Werff et al. 2009; Kariv et al. 2024), but a broadband masking noise is often still presented at ∼55-75 dB SPL especially if the results are inconclusive (Hatton et al. 2022 p.86; Ontario Ministry of Children and Youth Services 2020 pp.16, 53). In adults, however, the interaural attenuation is essentially nil, necessitating contralateral masking to ensure the tested ear drives the response. A recent study found that effective masking levels using white noise in adults are 65 dB SPL for 500 Hz tone pips at 20 dB nHL and 53 dB SPL for 2000 Hz tone pips at 30 dB nHL, with infants requiring similar masking for 2000 Hz but an additional 13 dB of masking at 500 Hz (Lau & Small 2021). But a broadband masker such as white noise may not be the most appropriate for pABR stimuli. The pABR stimuli are already more place-specific, providing inherent masking by the parallel/simultaneous stimulation (Stoll & Maddox 2023; Stoll & Maddox 2024). In this proof-of-concept study, we chose to use the pABR stimuli as a starting place for the contralateral masker but with 10 times the stimulus rate for more energy. Along with the ∼+20 dB from the faster stimulus rate, the same voltage outputs for the AC masker gave rise to ∼20–30 dB higher levels (Supplemental Digital Content, Supplemental Table 2), resulting in a masker +40 dB nHL higher than the BC stimulus at each level in the intensity series. With this masking, on average the contralateral response (excluding 8000 Hz) was 17–43 nV smaller and 0.1–1.2 ms later, although not statistically significant. This is consistent with previous work in adults, who show smaller ipsilateral/contralateral differences than infants (Foxe & Stapells 1993), who in turn will have up to ∼100 nV and 0.5–1 ms differences between channels (Foxe & Stapells 1993; Stapells & Ruben 1989). Thus, our patterns in adults suggest our masking may have been sufficient; however, other levels, stimulus rates, or stimuli could be more effective maskers for pABR BC. Future studies could systematically investigate the optimal pABR masking parameters for BC and for infants.

### Limitations & future directions

This study was a proof-of-concept for BC with parallel stimulation and thus has some limitations and need for future studies. As mentioned already, but summarized here, lower levels were tested with BC due to using the same voltage outputs driving the B-71 vibrator to a lower force level than for the ER-2 insert earphones. The reference values herein could be used to create stimuli with altered voltage outputs so that similar levels are presented through the vibrator and earphones during a pABR test session. We used the pABR stimuli with a higher stimulus rate with the same voltage outputs (thus higher levels) for contralateral AC masking, which overall evoked ipsilateral responses that were the same or larger size and the same or earlier latency, suggesting the ipsilateral ear was driving the BC response. However, this masking may not have been effective for every individual, and more systematic investigations could evaluate the most effective stimulus and masking levels, for both adults and for infants, who often require different masking levels (Lau & Small 2021). Next studies could also verify the accuracy of BC for estimating threshold, to verify eHL corrections for BC pABR stimuli.

### Conclusions

Whichever transducer was used, stimuli presented at the same perceptual level yield highly similar ABRs for bone and air conduction, indicating that bone conduction pABR is feasible. Bone conduction is an essential part of audiological diagnosis of hearing loss and works just as well with pABR as traditional serial methods. Together with establishing dB nHL values, this work represents an important step towards translating the pABR for diagnosing hearing loss in infants.

## Supporting information

Supplemental Digital Content

## Acknowledgments

This work was funded by the National Institutes of Health grant R01DC017962. Isabel Herb and Eric Mitchell helped with recruitment and data collection. Ben Eisenreich assisted with bone conduction calibration. Portions of this article were presented at the 52^nd^ Annual Scientific and Technology Conference of the American Auditory Society in February 2025, and Forum Acusticum EuroNoise in June 2025.

## References

Benjamini, Y., Hochberg, Y. (1995). Controlling the False Discovery Rate: A Practical and Powerful Approach to Multiple Testing. Journal of the Royal Statistical Society. Series B (Methodological), 57, 289–300.

Carhart, R., Jerger, J.F. (1959). Preferred Method For Clinical Determination Of Pure-Tone Thresholds. Journal of Speech and Hearing Disorders, 24, 330–345.

Foxe, J.J., Stapells, D.R. (1993). Normal infant and adult auditory brainstem responses to bone-conducted tones. Audiology, 32, 95–109.

Gorga, M.P., Kaminski, J.R., Beauchaine, K.L., et al. (1993). A Comparison of Auditory Brain Stem Response Thresholds and latencies Elicited by Air-and Bone-Conducted Stimuli. Ear and Hearing, 14, 85–94.

Gramfort, A., Luessi, M., Larson, E., et al. (2013). MEG and EEG data analysis with MNE-Python. Front. Neurosci., 7. Available at: https://www.frontiersin.org/articles/10.3389/fnins.2013.00267/full [Accessed July 22, 2019].

Green, P., MacLeod, C.J. (2016). SIMR: an R package for power analysis of generalized linear mixed models by simulation. Methods in Ecology and Evolution, 7, 493–498.

Hatton, J.L., Janssen, R.M., Stapells, D.R. (2012). Auditory Brainstem Responses to Bone-Conducted Brief Tones in Young Children with Conductive or Sensorineural Hearing Loss. Int J Otolaryngol, 2012, 284864.

Hatton, J.L., Van Maanen, A., Stapells, D.R. (2022). British Columbia Early Hearing Program: Auditory Brainstem Response (ABR) Protocol. Available at: http://www.phsa.ca/bc-early-hearing/Documents/ABR_Protocol.pdf.

Joint Committee on Infant Hearing (2019). Year 2019 Position Statement: Principles and Guidelines for Early Hearing Detection and Intervention Programs. Journal of Early Hearing Detection and Intervention, 4, 1–44.

Kariv, L., Taitelbaum-Swead, R., Levit, Y. (2024). Assessment of Interaural Attenuation in Infants and Young Children Using Bone-Conducted Auditory Brainstem Response. Ear Hear, 45, 999–1009.

Kuznetsova, A., Brockhoff, P.B., Christensen, R.H.B. (2017). lmerTest Package: Tests in Linear Mixed Effects Models. Journal of Statistical Software, 82, 1–26.

Larson, E., Gramfort, A., Engemann, D.A., et al. (2023). MNE-Python. Available at: https://zenodo.org/records/8322569 [Accessed October 6, 2025].

Larson, E., McCloy, D., Maddox, R., et al. (2014). expyfun: Python experimental paradigm functions, version 2.0.0. Available at: https://zenodo.org/record/11640/export/hx [Accessed August 6, 2020].

Lau, R., Small, S.A. (2021). Effective Masking Levels for Bone Conduction Auditory Brainstem Response Stimuli in Infants and Adults with Normal Hearing. Ear and Hearing, 42, 443.

Ontario Ministry of Children and Youth Services (2020). Protocol for Auditory Brainstem Response – Based Audiological Assessment (ABRA). Available at: https://www.uwo.ca/nca/pdfs/clinical_protocols/2018.02%20ABRA%20Protocol_Oct%202020.pdf.

Polonenko, M.J., Maddox, R.K. (2022). Optimizing Parameters for Using the Parallel Auditory Brainstem Response to Quickly Estimate Hearing Thresholds. Ear Hear, 43, 646–658.

Polonenko, M.J., Maddox, R.K. (2019). The Parallel Auditory Brainstem Response. Trends Hear, 23, 2331216519871395.

R Core Team (2025). R: A Language and Environment for Statistical Computing, Vienna, Austria: R Foundation for Statistical Computing. Available at: https://www.R-project.org/.

Sharma, M., Purdy, S.C., Bonnici, L. (2003). Behavioural and Electroacoustic Calibration of Air-conducted Click and Toneburst Auditory Brainstem Response Stimuli. Australian and New Zealand Journal of Audiology, 25, 54–60.

Stapells, D.R. (2011). Frequency-specific ABR and ASSR threshold assessment in young infants. Phonak Communications, 409–448.

Stapells, D.R. (2000). Threshold estimation by the tone-evoked auditory brainstem response: a literature meta-analysis. Canadian Journal of Speech-Language Pathology and Audiology, 24, 74–83.

Stapells, D.R., Oates, P. (1997). Estimation of the pure-tone audiogram by the auditory brainstem response: a review. Audiol Neurootol, 2, 257–280.

Stapells, D.R., Ruben, R.J. (1989). Auditory brain stem responses to bone-conducted tones in infants. Ann Otol Rhinol Laryngol, 98, 941–949.

Stoll, T.J., Maddox, R.K. (2023). Enhanced Place Specificity of the Parallel Auditory Brainstem Response: A Modeling Study. Trends in Hearing, 27, 23312165231205719.

Stoll, T.J., Maddox, R.K. (2024). Enhanced Place Specificity of the Parallel Auditory Brainstem Response: An Electrophysiological Study. J Assoc Res Otolaryngol, 25, 477–489.

Vallat, R. (2018). Pingouin: statistics in Python. Journal of Open Source Software, 3, 1026.

Vander Werff, K.R., Prieve, B.A., Georgantas, L.M. (2009). Infant air and bone conduction tone burst auditory brain stem responses for classification of hearing loss and the relationship to behavioral thresholds. Ear Hear, 30, 350–368.

