## Supplemental Digital Content for "Bone conducted responses using the parallel auditory brainstem response (pABR) paradigm"

### Supplemental Tables

Supplemental Table 1. Modeled correction factors (dB) for pABR stimuli.

| Tone pip | Stimulus Rate | | | | | | | | | | | |
|---|---|---|---|---|---|---|---|---|---|---|---|---|
| | 20 Hz | | | 40 Hz | | | 60 Hz | | | 80 Hz | | |
| | AC | BC | Both | AC | BC | Both | AC | BC | Both | AC | BC | Both |
| 500 Hz | 15.0 | 15.0 | 15.0 | 13.4 | 13.7 | 13.5 | 11.9 | 12.3 | 12.0 | 10.3 | 10.9 | 10.5 |
| 1000 Hz | 16.4 | 16.7 | 16.5 | 15.1 | 15.3 | 15.2 | 13.8 | 13.9 | 13.8 | 12.4 | 12.6 | 12.5 |
| 2000 Hz | 17.9 | 18.3 | 18.0 | 16.8 | 16.9 | 16.8 | 15.7 | 15.5 | 15.6 | 14.5 | 14.2 | 14.4 |
| 4000 Hz | 19.3 | 19.9 | 19.5 | 18.4 | 18.5 | 18.5 | 17.6 | 17.1 | 17.4 | 16.7 | 15.8 | 16.4 |
| 8000 Hz | 20.8 | 21.5 | 21.0 | 20.1 | 20.1 | 20.1 | 19.5 | 18.7 | 19.2 | 18.8 | 17.4 | 18.3 |

Note: AC = air conduction from full model, BC = bone conduction from full model, Both = common correction factor from parsimonious model

Supplemental Table 2. Level adjustments for the EEG data.

| | 500 Hz | 1000 Hz | 2000 Hz | 4000 Hz | 8000 Hz |
|---|---|---|---|---|---|
| A) Levels for the same voltage output of 70 dB* | | | | | |
| BC: B-71 (dB nHL) | 29.75 | 36.30 | 38.45 | 26.00 | 20.40 |
| AC: ER-2 (dB nHL) | 50.50 | 54.80 | 50.70 | 51.50 | 53.40 |
| Difference (dB) [AC – BC] | 20.75 | 18.50 | 12.25 | 25.50 | 33.00 |
| B) Level offsets for plotting waveforms in 10 dB steps | | | | | |
| Adjustment to BC waveforms (dB) | -20 | -20 | -10 | -20 | -30 |
| BC offset from AC (dB) in waveforms | -0.75 | 1.50 | -2.25 | -5.5 | -3.0 |
| C) Offsets from 10 dB steps to calculate actual levels (add to 10-dB step level) | | | | | |
| BC offset from 10 dB step-size (dB) | -0.25 | 6.30 | -1.55 | 6.00 | 0.40 |
| AC offset from 10 dB step-size (dB) | 0.50 | 4.80 | 0.70 | 1.50 | 3.40 |

* Calculated as measured dB (dyne or SPL) for pure tone – modeled correction factor – RETVFL or RETSPL

Note: AC = air conduction, BC = bone conduction, dB nHL = decibels normal hearing level, peSPL = peak-equivalent sound pressure level, EEG = electroencephalography

Supplemental Table 3. Linear mixed effects model of residual noise of the responses.

| Fixed Effect | Estimate | SE | df | t | p |
|---|---|---|---|---|---|
| Intercept (AC) | 23.932 | 2.391 | 458.726 | 10.011 | <0.001 |
| dBs | 0.035 | 0.118 | 753.000 | 0.295 | 0.768 |
| LogFreq | 0.296 | 0.662 | 753.000 | 0.448 | 0.655 |
| Trd(BC) | 1.732 | 3.142 | 753.000 | 0.551 | 0.581 |
| dBs:LogFreq | 0.001 | 0.035 | 753.000 | 0.020 | 0.984 |
| dBs:Trd(BC) | 0.085 | 0.177 | 753.000 | 0.483 | 0.629 |
| LogFreq:Trd(BC) | -0.464 | 0.946 | 753.000 | -0.491 | 0.624 |
| dBs:LogFreq:Trd(BC) | -0.037 | 0.054 | 753.000 | -0.697 | 0.486 |

Note: filtered levels within 0 to 30 dB nHL (inclusive).

LogFreq = log10(frequency in Hz), Trd = transducer, BC = bone conduction, default in the full model is air conduction (AC); SE = standard error

#### Supplemental Figures

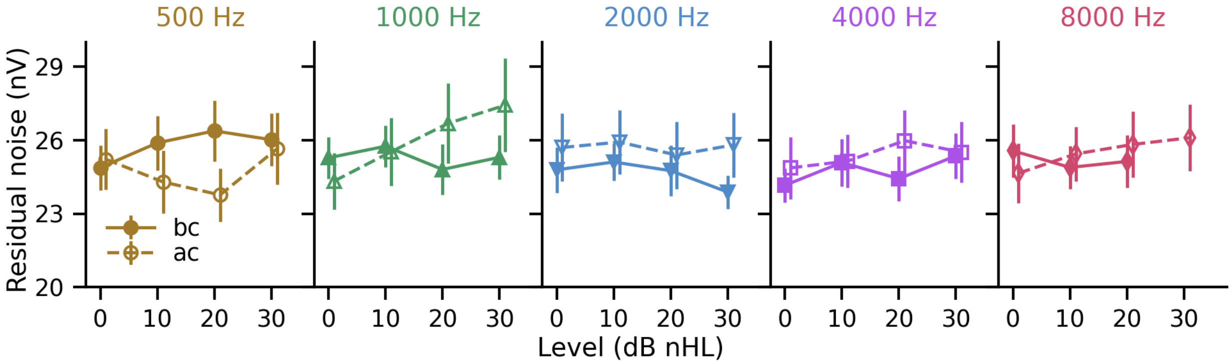


Supplemental Figure 1. Residual noise for air- (ac, open symbols and dashed line) and bone- (bc, closed symbols and solid line) conduction across frequency and level, calculated from -480 to -20 ms inclusive.
